# Decoding Protein Dynamics: ProFlex as a Linguistic Bridge in Normal Mode Analysis

**DOI:** 10.1101/2024.09.21.614246

**Authors:** Damian J. Magill, Timofey A. Skvortsov

## Abstract

Artificial intelligence has revolutionized structural bioinformatics, with AlphaFold being arguably the most impactful development to date. The structural atlases generated by these methods present significant opportunities for unraveling biological mysteries, but also pose challenges in leveraging such massive datasets effectively. In this work, we explore the dynamic landscape of hundreds of thousands of AlphaFold-predicted structures using normal mode analysis. The resulting data is used to define an alphabet summarizing relative protein flexibility, termed ProFlex. We believe that refining and further applying ProFlex-like approaches offers novel opportunities for understanding protein function and enhancing other methods.

## Introduction

Proteins are key building blocks of all living entities. Their functional activity is mainly determined by their structure and the knowledge of the structural organization of proteins is thus critically important for their better understanding and manipulation. Proteins are inherently flexible biological macromolecules and normal experimental methods of structural characterization of proteins such as X-ray do not capture conformational changes. Computational methods of structure prediction, including AlphaFold and similar AI-based approaches also do not include or use dynamic information. Protein conformational changes can be predicted with a number of approaches, including quantum mechanics/molecular mechanics (QM/MM) simulations, molecular dynamics (MD) simulations, and normal mode analysis (NMA).

Normal mode analysis is a method for characterizing the various flexible configurations that a protein or molecule can assume around a stable state. The concept revolves around the behavior of an oscillating system in equilibrium, such as a protein at its lowest energy shape. When this system is disturbed, it experiences a corrective force that aims to restore it to its original, stable shape, that exhibiting the least energy. Equilibrium for a system is defined as the point where all applied forces cancel out, resulting in no net force.

When we summarise this for an equilibrated system we get the following:

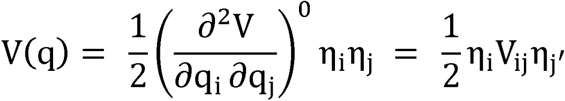

Here *qi*, and *qj* represent the current configuration of components i and j of the system and *V(q)* is the potential energy component of the system. *η_i_* represents the deviation of *i* from its equilibrium configuration. *V_ij_* is a hessian matrix that contains second order derivatives of the system potential (Goldstein, 1980).

This provides information on how components relate to one another when changes occur in the system. Of course, we also need to consider mass and kinetic energy in the determination of the final equation of motion. Despite this, by substituting an oscillatory equation into the equation of motion we can derive a standard eigenvalue equation as follows:

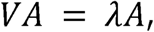

In summary, the matrix *A* contains eigenvectors *A_k_* from the Hessian matrix *V*, and *λ* is a diagonal matrix with eigenvalues *λ_k_.* The eigenvectors, or normal mode vectors, indicate the relative movement direction and magnitude for each particle in the system, while the eigenvalues represent the squared vibration frequencies of the particles in each mode (Bahar and Rader, 2005).

Normal modes offer analytical solutions to the equations of motion, predicting the future positions of atoms based on initial conditions, within the small oscillation approximation. Despite being much more computationally tractable compared to molecular dynamics simulations, coarse grained approaches, for example, the ones replacing semi-empirical potentials by simple harmonic potentials such as with the elastic network model, are often applied (Na et al. 2015).

The broader application of NMA has been proposed as a method to refine derived protein structures (Delarue, 2008; Suhre and Sanejouand, 2004). In the context of the extensive structural atlas generated by artificial intelligence advancements, such as AlphaFold2 and ESMFold, innovative approaches are required to fully leverage this data (Jumper et al. 2021; Rives et al. 2021). One such innovation is Foldseek, which utilizes the 3Di alphabet to convert 3D information (atom coordinates) into 1D (string of characters), enabling the use of conventional sequence search algorithms for comparison of structures while preserving most of the 3D information (Van Kempen et al 2024).

The concept of using alternative alphabets to represent various protein characteristics is well-established. For instance, hidden Markov model alphabet representations have been employed to encode protein folds and conserved domains into a single dimension (Deschavanne & Tufféry, 2009). Additionally, several studies have utilized reduced amino acid alphabets to simplify protein sequence complexity (Li et al., 2003; Petersen et al., 2009).

To date, large-scale applications of protein dynamics approaches have not been documented with existing studies targeting the protein structural universe in a static manner (Barrio-Hernandez et al., 2023). In this study, we present a large-scale NMA analysis of over 500,000 protein structures predicted by AlphaFold2. By deriving root mean square fluctuation (RMSF) values for all structures, we have empirically defined a specific alphabet called ProFlex, which facilitates the summarization and analysis of protein dynamic datasets. In addition to providing the means to apply sequence-based analysis methodologies for massive datasets, such approaches could additionally serve in refining structural alphabets that may struggle with highly flexible and disordered regions.

We provide an in-depth analysis of the ProFlex dataset in comparison to other protein information levels. Furthermore, we show that it is feasible to translate directly from amino acid sequences to the ProFlex alphabet using protein language models. Overall, we believe ProFlex- like approaches open new possibilities in structural bioinformatics that should be explored more deeply.

## Methods

### Dataset and Normal Mode Analysis

The dataset used for this study consisted of the SWISS-PROT AlphaFold 2 generated models (https://ftp.ebi.ac.uk/pub/databases/alphafold/latest/swissprot_pdb_v4.tar). Normal mode analysis and extraction of root mean square deviation (RMSF) values was conducted using the Bio3D package within the R programming environment (Grant et al. 2021). After the NMA analysis, files for which NMA analysis did not generate valid output were removed to retain only valid structures. The final dataset contained per-residue RMSF values in vectorized form for 526,820 protein models that were analysed in this study. The smallest and largest proteins in this dataset were 16 and 2699 amino acids respectively with an average size of 353 amino acids.

### Initial Evaluation of Dataset

To analyze the periodic patterns in the RMSF data, a Fast Fourier Transform (FFT) was applied to each RMSF vector using the FFT module in numpy (v2.1.1). The FFT converts the RMSF values from the time domain to the frequency domain, allowing the identification of dominant frequencies that characterize the flexibility profiles of the proteins. The magnitudes of the FFT coefficients were normalized using the StandardScaler from the Scikit-learn library (v1.5.2) to ensure comparability across different sequences. DBSCAN from the Scikit-learn library was employed to group the proteins based on their FFT features with the epsilon parameter determined via k-th nearest neighbours. Dimensionality reduction was then conducted using principal component analysis and t-distributed stochastic neighbor embedding. Euclidean distance was used in the extraction of centroids for each cluster.

### Determining ProFlex Alphabet

The RMSF data obtained was used to develop a protein alphabet we named ProFlex that can be used to encode the information about flexibility of each amino acid in a protein. A schematic overview of the data transformations to and from ProFlex is given in Fig 1.

#### Scaling Data and Determining Alphabet Size

Some proteins exhibit higher overall RMSF values compared to others, posing a challenge in defining a simple alphabet that captures the entire range of flexibility values while retaining adequate resolution to reflect changes in flexibility across a given structure. To address this, we employed min-max scaling across the entire dataset constrained between 0 and 1. Although this approach prevents direct comparison of flexibility magnitudes among structures, it allows us to retain information about relative flexibility within a given structure, which is our primary interest.

We aimed to define an alphabet that maximizes information capture while remaining relatively simple and easily exploitable. To determine the appropriate alphabet size and its underlying representation, an empirical approach was employed. We based our decision on global step sizes from our dataset, ensuring sufficient letters to capture changes in flexibility from one amino acid to the next. Our calculations revealed an average step size of 0.0226 scaled RMSF values, a median of 0.0156, and a mode of 0, indicating stretches of residues with the same flexibility. These values suggested a letter range of 46 to 67 to capture these specific step sizes. For simplicity, we chose to use the collective lower- and upper-case English alphabet, totaling 52 letters, which falls within this range.

With this alphabet and the modal step size of zero in mind, we plotted the overall step sizes of the dataset (Fig S1). The skew in step sizes suggests that equally binning values for the 0 to 1 range may not be the best approach. Therefore, we applied different binning methods to define the ranges of scaled RMSF values represented by the ProFlex alphabet.

#### Global Dynamic Binning

This approach seeks to divide the dataset into bins based upon the entire distribution. For a dataset *D* = {*d*_1_, *d*_2_, …, *d_N_*} where *N* is the total number of values bin determination can be defined as follows:

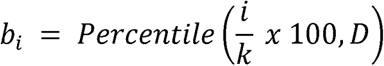

Where *k* corresponds to the number of bins in which we divide the dataset *D* in order to create *k + 1* bin edges for the *i*-th bin edge *b_i_*

Each value *d* ∈ *D* is mapped such that membership of the *i*-th bin is given by *b_i_*_−1_ ≤ *d* < *b_i_*

#### Sequence Specific Binning

This approach differs from the above in that the bin edges are defined by the value distribution of each sequence allowing for better per sequence bin adaptation. In this case we let *S_j_* = *S_j_*_1_, *S_j_*_2_, …, *S_jNj_* correspond to the set of RMSF for the *j*-th sequence where *N_j_* is the number of values in the *j*-th sequence. We then calculate percentiles for the sequence *S_j_* to define the bin edges. The *i*-th bin edge for sequence *j* is given as:

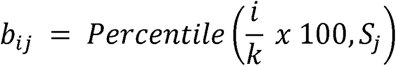

This creates *k* + 1 bin edge*s b_j_*_0_, *b_j_*_1_, …, *b_jk_* for each sequence *j* where *b_j_*_0_ is typically 0 and *b_jk_* is the maximum value in *S_j_*. Each value of *S_ji_* maps to the *m-th* bin *b_j,m−_*_1_ ≤ *S_j_* < *b_jm_* and subsequently to the *m*-th letter of the alphabet.

### Evaluating Alphabets

For each binning, bin edges were saved and used to map the alphabet back to scaled values, using the midpoint of each bin range. The resulting data distribution was statistically compared to the original scaled data using Wilcoxon tests for each sequence-reconversion pair. This was done to determine which binning approach deviates the least from the source data, considering the natural loss of resolution due to mapping back to midpoint bin values.

### Cluster Analysis of Alphabets

To directly compare datasets encoded as alphabets, a k-mer clustering approach was utilized. K- mer frequencies were established as a DictVectorizer object for k = 2 and 3 and standardized using min-max scaling (sci-kit learn v1.5.2). Optimal cluster numbers for each dataset were determined using the Elbow and Silhouette methods and then applied in k-means clustering (sci-kit learn v1.5.2). This was followed by dimensionality reduction and visualization of the two dominant principal components. To evaluate the structure of clusters across datasets for each k-mer value, the adjusted Rand index (ARI), normalized mutual information (NMI), homogeneity, completeness, and V-measure were all determined using Scikit-learn v1.5.2.

### Secondary structure analysis and 3Di Alphabet Determination

As an additional point of comparison secondary structure sequences were also determined for the entirety of the dataset. This was done using the DSSP utility from Bio.PDB (Kabsch and Sander., 1983). The subsequent output was simplified via the use of a secondary structure map that converted all variants of β-sheets to “E”, 310 and π helices to “H” along with conventional α helices, and finally all turns, bends, and coils were mapped to “C”.

In order to determine the 3Di alphabets for the dataset we used the mini3di package for Python (https://pypi.org/project/mini3di/#ref1).

### Molecular Modelling and Analysis

Molecular modeling of viral capsid proteins not included in the SWISS-PROT AF2 DB was conducted using the AlphaFold 2 package in monomer mode and employing the fully compiled database of PDB structures (Jumper et al. 2021). Predictions were scored based on the pLDDT criteria to choose the top ranked model. Visualization of models and preparation of publication grade figures was conducted within Pymol V2.4 (DeLano, 2002).

### Phylogenetic and Similarity Analysis

Sequence alignments were conducted using Clustal Omega and the neighbour joining method within MEGA10 leveraged for the reconstruction of phylogenetic relationships (Severs and Higgens, 2014; Kumar et al. 2018). In order to compare similarities between relationship inference at the amino acid, secondary structure, and ProFlex levels, all-vs-all sequence comparisons were conducted using a simple implementation of the Needleman-Wunsch algorithm to infer a distance matrix (Likic, 2008). To provide a comparison at the structural level, the TM-Align algorithm was used and the resulting scores also compiled to form a distance matrix (Zhang and Skolnick., 2005).

### Translating from Amino Acids to ProFlex alphabet

In order to evaluate the feasibility to translate directly from amino acid sequences to the ProFlex alphabet we implemented a transformer encoder-decoder architecture for a subset of sequences lying in the 150 – 300 amino acid range. This range was chosen both to eliminate outliers from the NMA analysis and to conform to computational constraints.

Character level tokenization was followed by embedding to 128 dimensional vectors boosted with positional embeddings. Transformer layers comprised feed forward networks possessing 2048 intermediate dimensionality with 8 multi-attention heads. The decoder architecture mirrors this with the application of softmax activation (Fig 2). A custom loss function was applied based on sparse categorical entropy with incorporated masking for padding tokens. Adam optimization was applied with initial learning rate of 0.001 and a dropout rate of 0.1 employed. Validation loss and accuracy were monitored continually with early stopping applied to prevent overfitting prior to training for 300 epochs.

The trained model file and scripts are available at https://github.com/DamianJM/proFlex

### ProFlex Tool Suite

#### Construction of ProFlex Search Databases

In order to provide a rapid means of constructing and querying ProFlex alphabets, we opted to implement an n-gram based approach. Following the calculation of n-grams within protein sequences, a database is constructed by indexing each n-gram with the protein sequence IDs in which it appears for a given set of sequences.

Queries are subsequently divided into n-grams and searched against the database for which sequence IDs are retrieved for those sequences containing a specific n-gram from the query sequence. This process is repeated for all subsequences of the query and cumulative scores tallied. We then rank the protein sequences based on their scores and retrieve the top (*X*) sequences. Assuming Score (Si, Q) to be the cumulative score for protein sequence *Si*, based on n-grams matching the query. The top *X* protein sequences are those that maximize:

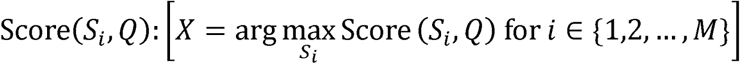

This approach results in a time complexity of *0*(*LQ*+ *MlogM*) due to the requirement to extract n-grams from a variable length query sequence LQ and *MlogM* corresponding to the time needed to sort alignment scores for a database containing *M* sequences.

#### ProFlex Suite Features

As part of this work we developed a suite of tools that allow users to easily leverage ProFlex in local workflows. Available as a series of importable modules along with a pre-compiled version of the SWISS-PROT dataset used in this study, users can directly query PDB files against local databases. The empirically determined bins for ProFlex determination are included along with the C-alpha NMA setup used for fast PDB to ProFlex conversion. This outputs an easy to interpret report in HTML format displaying the top 5 ProFlex hits along with amino acid and structural alignments for the best hit. Additionally, query and top hit ProFlex sequences are back translated and provided as a comparative flexibility profile graph (Fig S2).

The ProFlex Suite modules and guide on how to implement into local pipelines is available at https://github.com/DamianJM/proFlex

## Results and Discussion

### Global Analysis of RMSF Profiles

Before analyzing the ProFlex dataset, we examined the raw RMSF profile (vectors) dataset to gain insights into the global flexibility profiles. We used Fourier analysis to identify changes in flexibility across the protein structures. Clustering this data provided a global overview of the different flexibility profiles (Fig 2).

Approximately 51% (250k) of the full dataset (500k) was classified as “noise” by the algorithm, indicating a highly diverse data distribution that makes grouping based solely on raw flexibility values challenging. Surprisingly, the remaining data formed only 28 clusters, with one cluster comprising 48% (∼249k) of the remaining dataset. The centroid of this cluster was identified as a penicillin-binding protein 2B from *Streptococcus pneumoniae*. The other clusters were outliers, each containing between 2 and 19 members.

Plotting the RMSF values for the centroids revealed distinct periodicity in the profiles (Fig 2B), operating on different RMSF scales, which explains why it was detectable by Fourier analysis. However, this periodicity is an assumption and may not reflect actual protein dynamics, representing a significant limitation of this analysis and one that ProFlex seeks to address.

### ProFlex Alphabet Derivation

To determine the optimal approach for ProFlex alphabet derivation, we statistically compared three different binning methods to initial scaled RMSF values (Fig 3). The statistical analysis of the equal binning approach revealed significant shortcomings, with only one p-value above the 0.001 threshold. It was necessary to lower this threshold by three orders of magnitude to obtain just 3,384 values (0.64% of the dataset).

The differences between the binning approaches are visually evident in the example plots for randomly selected sequences. For the global binning approach, we observed 430,778 (81.8%), 485,030 (92.1%), and 512,078 (97.2%) values above the p-value thresholds of 0.05, 0.01, and 0.001, respectively. The sequence-based approach yielded higher values: 458,077 (86.9%), 498,714 (94.7%), and 516,437 (98.0%) for the same thresholds. Although the sequence-based approach performed better, the differences were surprisingly small, indicating that the global dataset’s bin edges are highly representative of most sequences. This is evident from the plots of both reconverted sequences of RMSF profile of randomly extracted proteins (Fig 3 bottom).

Given the marginal improvement offered by sequence-based binning, we opted for the global binning approach. Sequence-based binning and resulting alphabets may not be directly comparable between sequences, as some letters could map to different bin ranges in different sequences, limiting further analyses. Additionally, the sequence-based approach showed a peak of low p-values and high p-values, indicating it produced identical distributions in some cases but struggled in others, especially with larger proteins.

### Global Alphabet Comparisons and Cluster Analysis

The global datasets, comprising raw RMSF data, secondary structure predictions, and ProFlex alphabets, were analyzed, yielding several interesting insights. The distribution of amino acids in relation to both secondary structure and ProFlex alphabets was determined (Fig 4).

Overall, leucine was the amino acid most frequently found within alpha helix components, with 54% of leucine residues located in these regions. This aligns with leucine’s characteristics as a small hydrophobic amino acid. Valine residues were predominantly observed in sheet regions, accounting for 35%, likely due to the stabilizing hydrophobic contributions from its branched side chain. Proline was a dominant component of coiled regions, with 73% of proline residues found in these areas, consistent with its tendency to induce kinks in structures.

Comparing the amino acid distribution to the ProFlex alphabet revealed a relatively equal distribution across all residues. This is not surprising, as flexibility is a product of the entire structure, allowing residues in both locally flexible and rigid regions to exhibit large-scale motions due to the global structure. The only outlier was a 7% proportion of ProFlex “Z” attributed to methionine residues, likely due to methionine’s presence at the N-terminus, which is often a flexible component of the structure.

A final comparison of ProFlex versus secondary structure elements showed a clear trend. Less flexible letters of the ProFlex alphabet were predominantly found within defined secondary structure elements, with the first letters “a” and “b” showing the highest proportions of sheet and helix inclusion at 32% and 46%, respectively. More flexible letters were associated with a greater coil component, with the final letter “Z” found in 76% of such regions.

Sequence-based clustering for equivalent k-mer sizes (k=2 and k=3) revealed substantial differences in cluster structures across all alphabets. To derive meaningful insights from these differences, it is necessary to scale cluster comparison metrics (Fig. 5, Fig. S3, Fig. S4). This underscores the significant variation in information provided by each alphabet, which is crucial for studies of this magnitude.

When comparing clusters across alphabets, it is noteworthy that for both k-mer sizes, the highest agreement across all metrics is observed when comparing amino acid-based clustering to secondary structure alphabet clustering. Specifically, 3Di clustering exhibited higher homogeneity and Adjusted Rand Index (ARI) scores with ProFlex for both 2-mer and 3-mer based clustering, indicating higher cluster purity for this pair compared to 3Di and other alphabets. Similar trends were observed for ProFlex compared to secondary structure clustering, with the former showing substantially lower similarity with amino acid-defined clusters. This could be expected, as ProFlex is derived directly from structural information and is thus positioned further in information space from the primary sequence.

When comparing clustering results for the same alphabet at the 2-mer and 3-mer levels, both secondary structure and ProFlex alphabets demonstrated good agreement, while other alphabets showed substantial differences despite their structural diversity. This is likely due to the presence of homopolymeric regions in ProFlex and secondary structure alphabets, leading to equivalent k-mer abundances at the 2-mer and 3-mer levels.

In summary, global clustering results exhibit significant differences in all cases and should be applied on a case-by-case basis depending on the specific requirements.

### Analysis of ProFlex and NMA Dataset

Analysis of the global ProFlex alphabet shows that all states have relatively equal occupancy, except for the most flexible letter “Z.” This is because the alphabet is a relative flexibility metric, and all proteins will have at least one “Z.”

An analysis of the raw average RMSF values in relation to sequence size (Fig 6A) revealed a general trend of lower overall flexibility as sequence size increases. This trend is not explained by the distribution of secondary structure elements with size, indicating that it is a product of the global nature of the protein rather than more of the structure being found within defined secondary structure elements (Fig 6B).

Interestingly, both the most and least flexible proteins were among the smallest. Observations of the underlying structure revealed these to be small, highly ordered, and unordered peptides, respectively (Fig 6A), explaining the underlying dynamics.

To understand how flexibility changes across given structures, we established a transition matrix for the ProFlex alphabet (Fig 6C). This matrix summarizes future ProFlex states in relation to past ones, revealing general flexibility trends. Overall, there is a clear pattern where a given ProFlex state is typically followed by the same or a similar one, mirroring step size patterns previously observed in the raw NMA data and supporting homopolymeric regions in ProFlex representations. This is especially true for the least and most flexible components of the proteins. As we move towards more centrally located letters, we observe a slightly more diffuse distribution with increased transitions into different but nonetheless similar flexibility states. This likely represents local fluctuations in flexibility captured by the global dynamic binning approach.

### Information Richness of Protein Alphabets

Each alphabet provided here represents a different level of protein information in varying detail. To compare these alphabets, we assess their information richness by examining the ratio of theoretical to empirical Shannon Entropy. The ratios for the alphabets 3di, SS, ProFlex, and amino acids are 0.86, 0.94, 0.67, and 0.96, respectively. This indicates that amino acids and secondary structure alphabets fully leverage their potential for information representation. In contrast, 3di and ProFlex show lower scores, with 3di having the same theoretical entropy as amino acids. ProFlex, despite its high theoretical richness due to its complexity, shows higher empirical richness than 3di, suggesting a greater distribution in letter usage. The lower ratios in both cases highlight potential biased symbol usage, possibly linked to biological constraints at the structural level.

When examining k-mer frequency distributions for k up to 5 (Figure S5), we observe high occupancy for top k-mers in the secondary structure alphabet, which is expected. However, there is also high occupancy for two k-mers in the 3di alphabet, indicating many residues share the same local residue neighborhood. Amino acids and ProFlex display a flat representation overall, with ProFlex showing a slight elevation in the 4-mer and 5-mer ranges, indicating balanced frequency for these top k-mers. Comparing the ratio of theoretical to observed k-mers, amino acids dominate, with almost all possible combinations existing even at the 5-mer level. Structural alphabets show lower ratios, reflecting specific biological constraints in 3D space. ProFlex exhibits the lowest ratio, largely due to its higher complexity, resulting in more theoretical k-mers. The natural bias towards the most and least flexible letters in ProFlex, combined with coordinated motions described by NMA analysis, makes the observance of specific k-mers, such as ZaZaZa, virtually impossible. While a complex alphabet is necessary to represent specific flexibility ranges adequately, the low abundance of observed k-mers compared to amino acids suggests the potential utility of a simpler representation, where collective RMSF values for amino acid k-mers are mapped to single ProFlex letters.

### ProFlex as a Phylogenetic Tool

Recent interest has surged in using protein models to reconstruct phylogenetic relationships (Mifsud et al., 2024). This approach offers the advantage of resolving distant relationships that might be obscured by frequent sequence substitutions. However, direct structural comparisons are not only computationally intensive but also potentially limited by slight conformational changes affecting rigid body comparisons. Therefore, there is interest in reducing this to a 1D problem, such as with Foldseek (Moi et al. 2023), and in finding ways to incorporate and combine information at various levels depending on the phylogenetic problem.

We analyzed 15 major capsid protein sequences from a series of highly diverse *Tevenvirinae* phages, which are somewhat challenging to classify. We calculated similarity matrices based on amino acid sequences, secondary structure alphabets, 3Di alphabets, structural comparisons using TM-Align, and the ProFlex alphabet (Fig 7). Across all measures, we identified similarities in characterized clusters, demonstrating that all methods can capture underlying relationships, albeit with some notable differences.

When comparing relationships established from amino acid sequences and structural comparisons, we observed greater similarity in defined clusters, highlighting the higher level of structural conservation despite sequence divergence. However, there are substantial differences to note. For instance, comparing T4 to pf16, we observed an amino acid identity of only 38% but a TM-Align score of 0.9 (Fig 7). In contrast, amino acid comparisons of T4 with Shigella phage Sf24 showed 97.89% similarity but a lower TM-Align score of 0.79. Structural observations indicate that these differences are due to different conformations (Fig 7).

Similarity scores from secondary structure alphabets appear to offer a good compromise between amino acid sequences and structural alignments. This method captures close relationships shown at the amino acid level, such as with T4 and Sf24, while retaining more distant relationships observed with pf16. Some clusters with high structural conservation are closer to their amino acid sequence counterparts in this representation. However, it is important to note that the global similarity score scale is lower, indicating lower overall distances compared to amino acid sequences.

Moving to the 3Di comparisons we see mixed elements from the structural, amino acid, and secondary structure alphabets with the latter being the closest overall. It appears this alphabet offers a good compromise of all information in this particular case.

The ProFlex comparisons showed the most divergent set of relationships. Highly conserved clusters at the structural level also showed conservation at the ProFlex level but with greater overall dissimilarity. This was evident in examples that were similar in structural comparisons but not at the amino acid level. Conversely, some relationships deemed distant at the ProFlex level were captured by other methods, such as the similarity between T4 and Citrobacter phage Moon. In cases where structural similarity is lower compared to amino acid similarity, the ProFlex level shows even greater distance.

An appropriate combination of static structural information and dynamic ProFlex-encoded information may yield interesting results. Overall, it is clear that the different methods provide varying degrees of information. Indeed, a critical point is that whilst each method shares the underlying relationship it does not do so with the same level of similarity. The intelligent incorporation of all these methods could be valuable in addressing diverse phylogenetic questions and should be explored more.

### From amino acids to ProFlex?

The properties of a protein are determined by its amino acid sequence. We aimed to explore the feasibility of translating amino acids directly into ProFlex using natural language processing (NLP) techniques. Our translation model achieved an accuracy of 0.7178 on the validation set. Although this is not exceptional, it suggests that such approaches are feasible.

We tested a random sequence excluded from the model, a 150-amino-acid protein annotated as a ribosome maturation factor. This sequence was analyzed using NMA, and the scaled RMSF values were derived using globally defined percentiles. When graphed (Fig 8), the predicted flexibility closely matched the original values, despite some differences in flexible regions. However, the model struggled with robust predictions in other examples and requires improvement.

By leveraging larger pre-trained architectures, the vast number of available predicted structures, and possibly a refined alphabet, we believe this approach can be enhanced. This work serves as a proof of concept and highlights the potential for further exploration of protein dynamics. We hope it will inspire interest in bridging the sequence-functionality gap, particularly by refining 3Di-like alphabets to better account for disordered regions, where determining nearest neighbors in three-dimensional space is challenging.

## Conclusion

In this study, we conducted a large-scale normal mode analysis of AlphaFold-predicted protein structures, introducing the ProFlex alphabet as a novel method to summarize protein flexibility. Our findings demonstrate that ProFlex can effectively capture and represent the dynamic landscape of protein structures, providing a valuable tool for downstream analyses such as sequence-based searches and structural refinement.

Future work should focus on refining the ProFlex methodology and exploring its applications in various domains of structural bioinformatics. The integration of ProFlex with other structural and sequence-based tools holds promise for advancing our understanding of protein behavior and enhancing the accuracy of structural predictions. This could take the form of ProFlex based structural validation wherein unexpected sequence to structure deviations may indicate incorrect predictions of the latter.

Overall, ProFlex represents a significant step forward in the field of protein dynamics, offering new opportunities for research and discovery in structural bioinformatics.

---

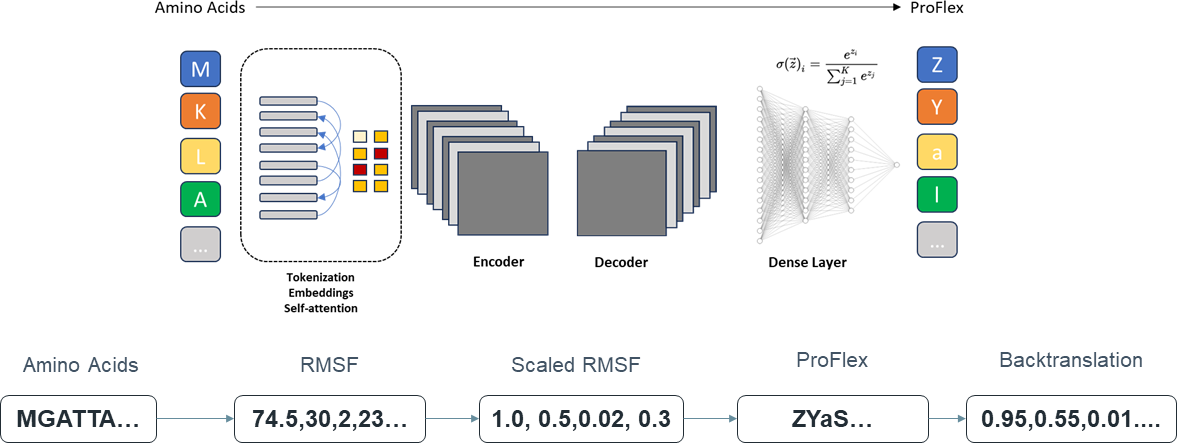

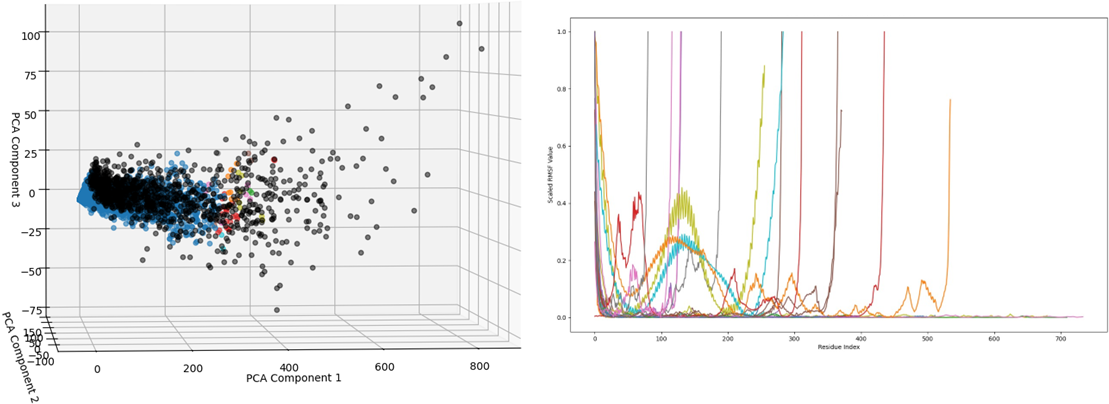

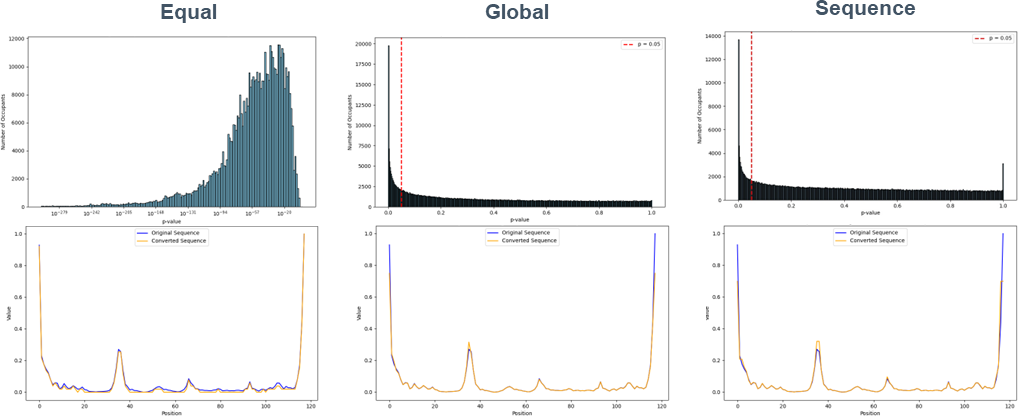

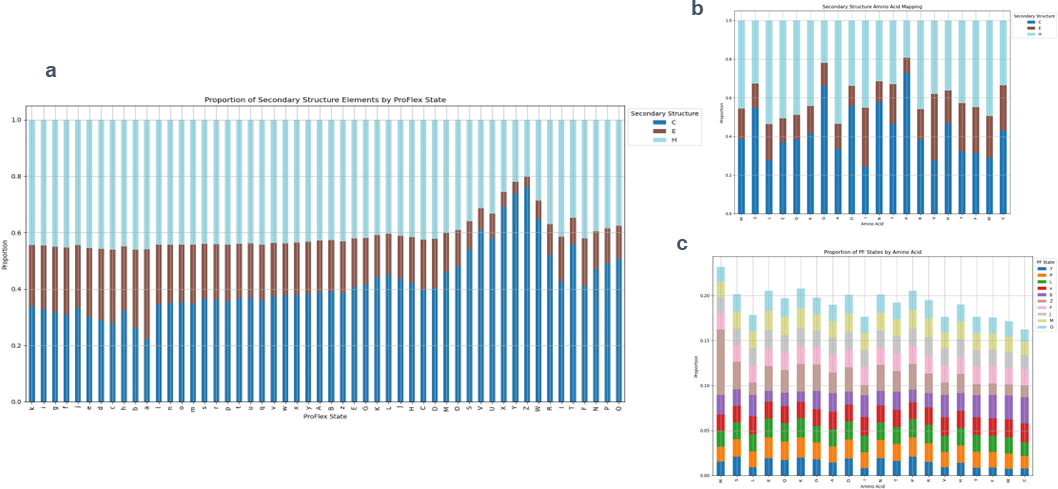

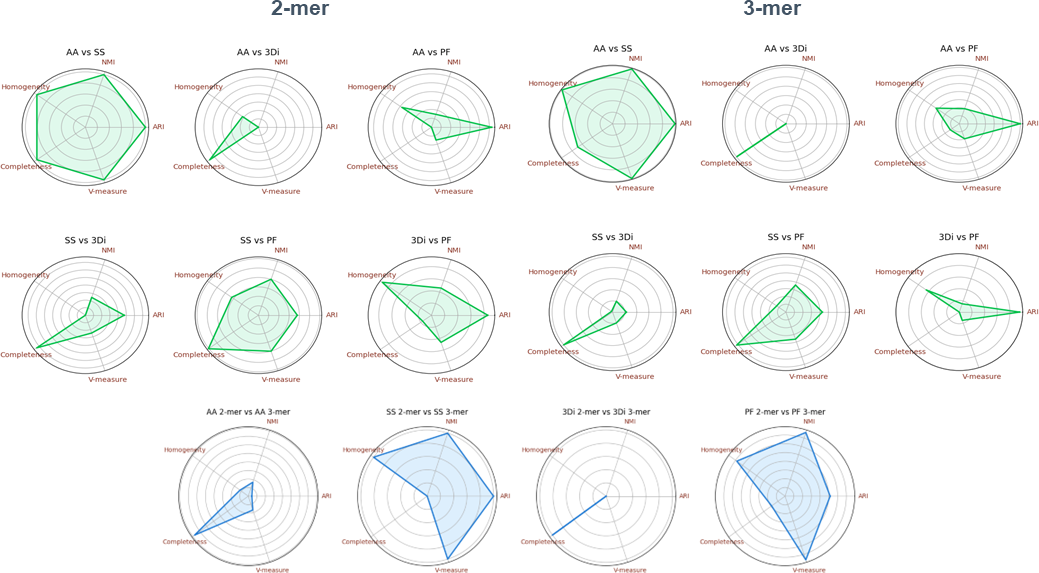

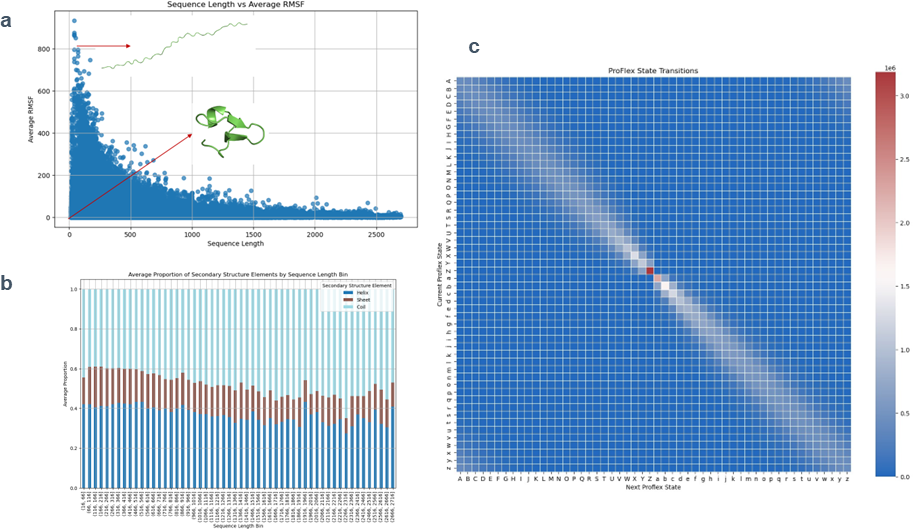

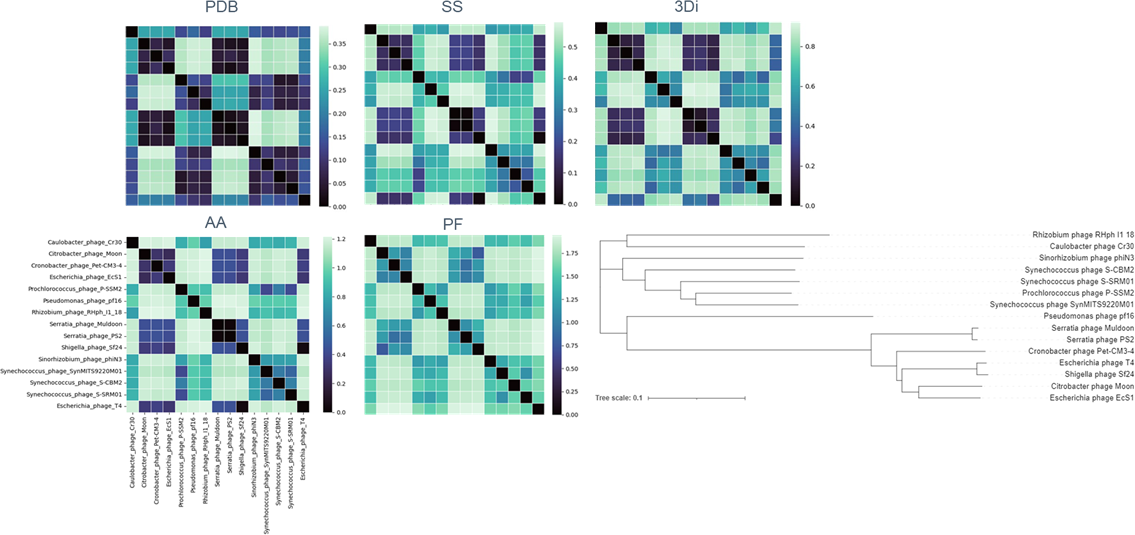

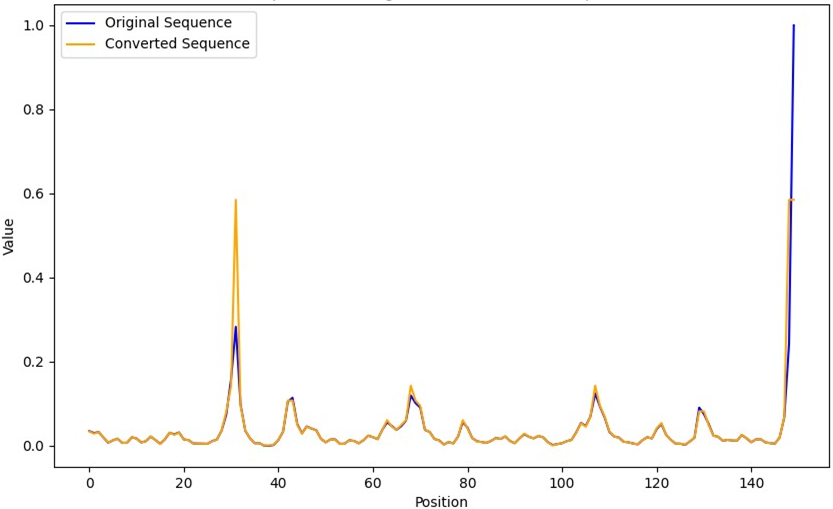

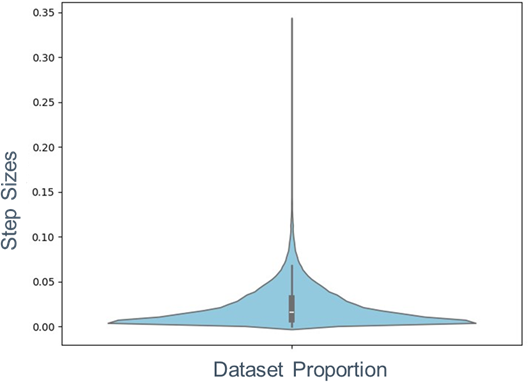

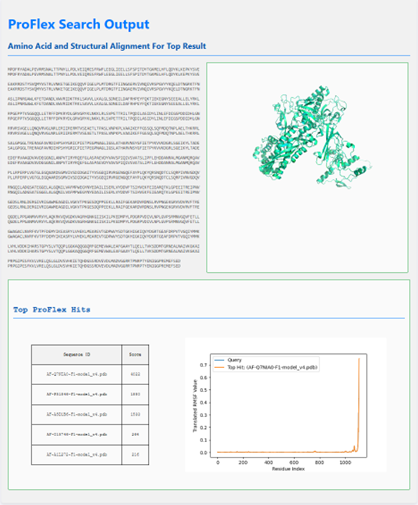

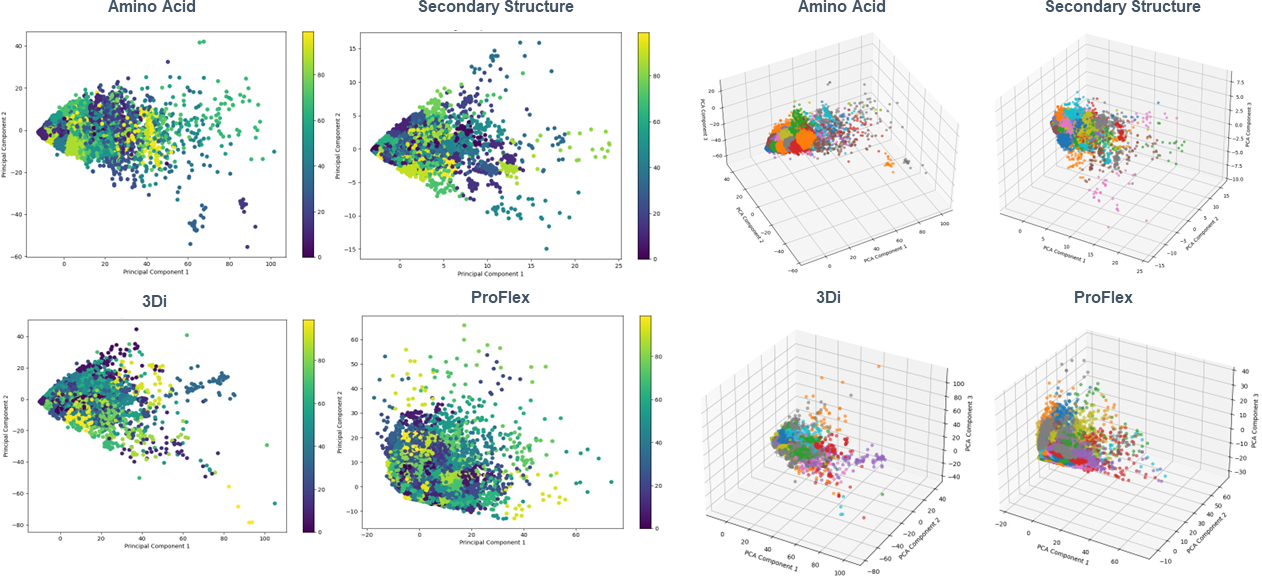

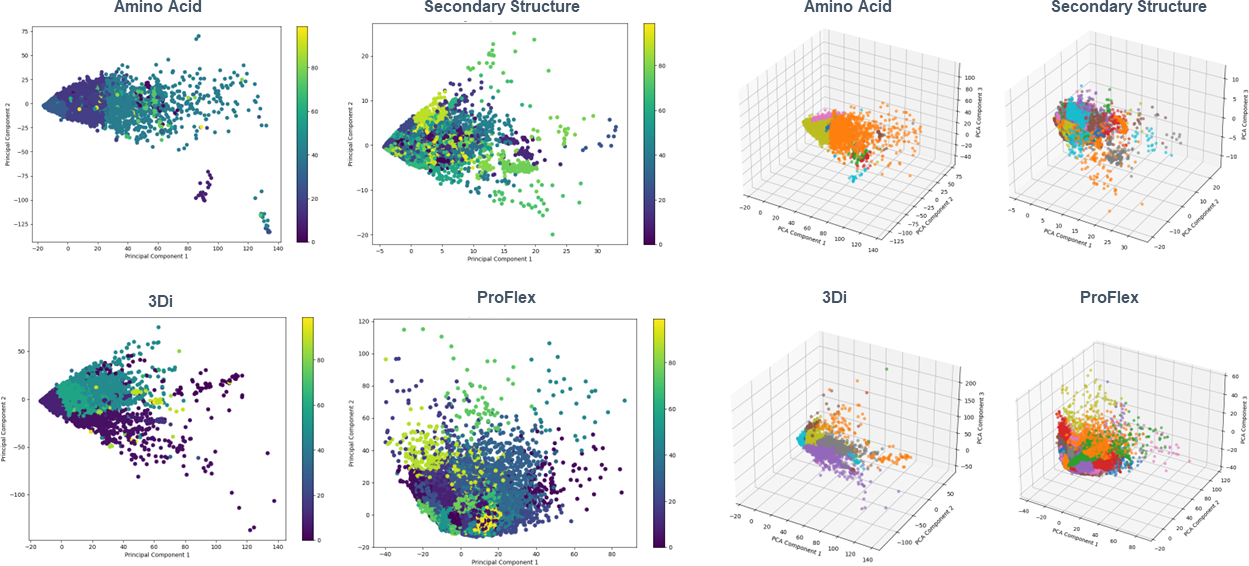

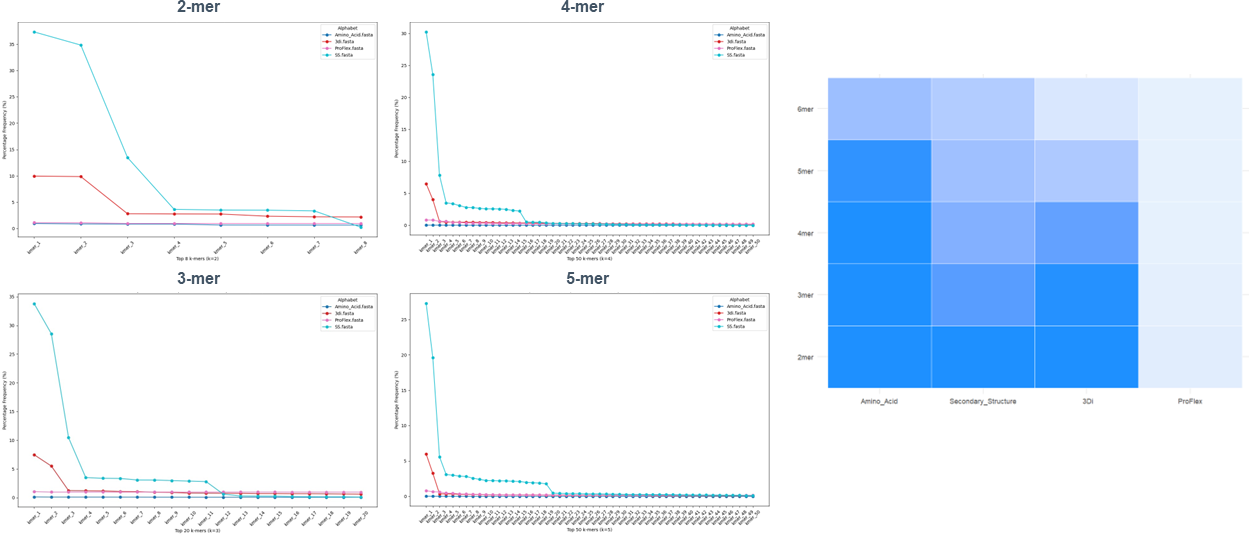

## References

Bahar, I. and Rader, A.J., 2005. Coarse-grained normal mode analysis in structural biology. Current opinion in structural biology, 15(5), pp.586–592.

Barrio-Hernandez, I., Yeo, J., Jänes, J., Mirdita, M., Gilchrist, C.L., Wein, T., Varadi, M., Velankar, S., Beltrao, P. and Steinegger, M., 2023. Clustering predicted structures at the scale of the known protein universe. Nature, 622(7983), pp.637–645.

DeLano, W.L., 2002. Pymol: An open-source molecular graphics tool. CCP4 Newsl. Protein Crystallogr, 40(1), pp.82–92.

Delarue, M., 2008. Dealing with structural variability in molecular replacement and crystallographic refinement through normal-mode analysis. Acta Crystallographica Section D: Biological Crystallography, 64(1), pp.40–48.

Deschavanne, P. and Tufféry, P., 2009. Enhanced protein fold recognition using a structural alphabet. Proteins: Structure, Function, and Bioinformatics, 76(1), pp.129–137.

Li, T., Fan, K., Wang, J. and Wang, W., 2003. Reduction of protein sequence complexity by residue grouping. Protein Engineering, 16(5), pp.323–330.

Grant, B.J., Skjærven, L. and Yao, X.Q., 2021. The Bio3D packages for structural bioinformatics. Protein Science, 30(1), pp.20–30.

Goldstein, H. Classical Mechanics, 2nd ed.; Addison-Wesley: Reading, MA, USA, 1980; pp. 243–274.

Jumper, J., Evans, R., Pritzel, A., Green, T., Figurnov, M., Ronneberger, O., Tunyasuvunakool, K., Bates, R., Žídek, A., Potapenko, A. and Bridgland, A., 2021. Highly accurate protein structure prediction with AlphaFold. nature, 596(7873), pp.583–589.

Kabsch, W. and Sander, C., 1983. Dictionary of protein secondary structure: pattern recognition of hydrogen-bonded and geometrical features. Biopolymers: Original Research on Biomolecules, 22(12), pp.2577–2637.

Kumar, S., Stecher, G., Li, M., Knyaz, C. and Tamura, K., 2018. MEGA X: molecular evolutionary genetics analysis across computing platforms. Molecular biology and evolution, 35(6), pp.1547–1549.

Likic, V., 2008. The Needleman-Wunsch algorithm for sequence alignment. Lecture given at the 7th Melbourne Bioinformatics Course, Bi021 Molecular Science and Biotechnology Institute, University of Melbourne, pp.1–46.

Mifsud, J.C.O., Lytras, S., Oliver, M.R. et al. 2024. Mapping glycoprotein structure reveals Flaviviridae evolutionary history. Nature 633, 695–703.

Moi, D., Bernard, C., Steinegger, M., Nevers, Y., Langleib, M. and Dessimoz, C., 2023. Structural phylogenetics unravels the evolutionary diversification of communication systems in gram- positive bacteria and their viruses. BioRXiv, pp.2023–09.

Na, H., Jernigan, R.L. and Song, G., 2015. Bridging between nma and elastic network models: preserving all-atom accuracy in coarse-grained models. PLoS computational biology, 11(10), p.e1004542.

Peterson, E.L., Kondev, J., Theriot, J.A. and Phillips, R., 2009. Reduced amino acid alphabets exhibit an improved sensitivity and selectivity in fold assignment. Bioinformatics, 25(11), pp.1356–1362.

Rives, A., Meier, J., Sercu, T., Goyal, S., Lin, Z., Liu, J., Guo, D., Ott, M., Zitnick, C.L., Ma, J. and Fergus, R., 2021. Biological structure and function emerge from scaling unsupervised learning to 250 million protein sequences. Proceedings of the National Academy of Sciences, 118(15), p.e2016239118.

Sievers, F. and Higgins, D.G., 2014. Clustal omega. Current protocols in bioinformatics, 48(1), pp.3–13.

Suhre, K. and Sanejouand, Y.H., 2004. On the potential of normal-mode analysis for solving difficult molecular-replacement problems. Acta Crystallographica Section D: Biological Crystallography, 60(4), pp.796–799.

Van Kempen, M., Kim, S.S., Tumescheit, C., Mirdita, M., Lee, J., Gilchrist, C.L., Söding, J. and Steinegger, M., 2024. Fast and accurate protein structure search with Foldseek. Nature biotechnology, 42(2), pp.243–246.

Zhang, Y. and Skolnick, J., 2005. TM-align: a protein structure alignment algorithm based on the TM-score. Nucleic acids research, 33(7), pp.2302–2309.

