## Supplementary material for "Decoding Protein Dynamics: ProFlex as a Linguistic Bridge in Normal Mode Analysis": Legends

**Fig 1. Conceptual overview of the transformer architecture used for amino acid to proflex translation (top)** **and overview of transformation process from amino acids to ProFlex (bottom)**

**Fig 2. Fourier analysis of Global NMA Derived RMSF Dataset**

3D PCA of DBSCAN clusters highlighting a huge flexibility space and lack of periodicity in protein flexibility (left) and scaled RMSF plots of cluster centroids showing examples of proteins exhibiting periodicity in their flexibility profiles (right)

**Fig 3. Comparison of equal, global, and sequence specific binning approaches to ProFlex alphabet determination.** Proflex alphabets were back translated to scaled RMSF values using mid-point percentile values and compared to original values using Wilcoxon tests (top). Examples of RMSF curves for original and backtranslated values are given for each binning approach (bottom).

**Fig 4. Global comparisons of amino acid, secondary structure, and ProFlex alphabets**

Proportion of proflex states found within specific secondary structure elements (a), secondary structure content divided by amino acids (b), and subset of proflex states divided by amino acids (c)

**Fig 5. Cluster comparisons from k-mer based clustering of different alphabets**

Comparisons for 2-mer and 3-mer levels provided across adjusted rand index, normalized mutual information, v-measure, completeness, and homogeneity for clusters generated for all alphabets.

**Fig 6. Global trends in the dataset with increasing sequence size**

Plot of average RMSF value against sequence size with example structures for least and most flexible representatives (a), distribution of secondary structure elements with increasing sequence size (b), and matrix of proflex transition states highlighting local flexibility changes in sequence space.

**Fig 7. Phylogenetic and similarity analysis of selected *Tevenvirinae* major capsid protein sequences and structures**

Similarity matrices for all alphabets have been generated on the basis of all-vs-all needleman Wunsch global alignment scores with the exception of structural comparisons (PDB) which are interpreted from inverted TM-Align scores. Phylogenetic trees were constructed on the basis of amino acid alignment

**Fig 8. Example of backtranslated scaled RMSF distribution of amino acid translated proflex sequence (orange) compared to original sequence (blue)**

**Fig S1. Violin plot of min-max scaled root mean square deviation step sizes used to determine appropriate ProFlex alphabet size**

**Fig S2. Example output provided by the ProFlex Tool Suite**

From a query PDB file users are provided with top proflex hits along with amino acid and structural alignments

**Fig S3. 2D and 3D PCA Plots for 2-mer abundance-based clustering**

**Fig S4. 3D PCA Plots for 2-mer abundance-based clustering**

**Fig S5. Information richness of alphabets as determined through kmer frequency distributions**

Most abundant kmers are presented for all alphabets for k ranging from 2 to 5 with abundance given as percentage in all instances. Heatmap displays proportion of actual to theoretical kmers for each alphabet for the SWISS-PROT data set
